# Nitrate Assimilation Underlying Kleptoplasty

**DOI:** 10.1101/2022.12.04.518691

**Authors:** Moe Maruyama, Tsuyoshi Kagamoto, Yuga Matsumoto, Ryo Onuma, Shin-ya Miyagishima, Goro Tanifuji, Masami Nakazawa, Yuichiro Kashiyama

**Affiliations:** Graduate School of Engineering, Fukui University of Technology, Fukui, 910-8505, Japan; Department of Applied Chemistry and Food Science, Fukui University of Technology, Fukui, 910-8505, Japan; Department of Gene Function and Phenomics, National Institute of Genetics, Mishima, 411-8540, Japan; Kobe University Research Center for Inland Seas, Awaji, 656-2401, Japan; National Museum of Nature and Science, 4-1-1, Amakubo, Tsukuba, 305-0005, Japan; Department of Applied Biochemistry, Faculty of Agriculture, Osaka Metropolitan University, Sakai, 599-8531, Japan

**Keywords:** chloroplast evolution, Euglenophyceae, horizontal gene transfer, kleptoplasty, nitrate assimilation, *Rapaza viridis*

## Abstract

While photoautotrophic organisms utilize inorganic nitrogen as the nitrogen source, heterotrophic organisms utilize organic nitrogen and thus do not generally have an inorganic nitrogen assimilation pathway. Here we focused on the nitrogen metabolism of *Rapaza viridis*, a unicellular eukaryote exhibiting kleptoplasty. Although belonging to the lineage of essentially heterotrophic flagellates, *R. viridis* exploits the photosynthetic products of the kleptoplasts and was therefore suspected to potentially utilize inorganic nitrogen. From the transcriptome data of *R. viridis*, we identified the gene Rv*NaRL*, which had sequence similarity to nitrate reductases found in plants. Phylogenetic analysis revealed that Rv*NaRL* was acquired by a horizontal gene transfer event. To verify its function of the protein product *Rv*NaRL, we established a RNAi mediated knockdown and a CRISPR-Cas9-mediated knockout experiments for the first time in *R. viridis* and applied them to this gene. The Rv*NaRL* knockdown and knockout cells exhibited significant growth only when ammonium was supplied but, in contrast to the wild-type cells, no substantial growth when nitrate was supplied. Such arrested growth in absence of ammonium was attributed to impaired amino acid synthesis due to the deficiency of nitrogen supply from the nitrate assimilation pathway; this in turn resulted in the accumulation of excess photosynthetic products in the form of cytosolic polysaccharide grains as observed. These results indicate that *Rv*NaRL is certainly involved in nitrate assimilation by *R. viridis*. Thus, we infer that *R. viridis* achieved its advanced kleptoplastic strategy owing to a posteriori acquisition of the nitrate assimilation pathway the horizontal gene transfer.

## Introduction

Photoautotrophic organisms acquire the nitrogen required for their biomolecules by assimilating inorganic nitrogen. Although nitrogen assimilation is ultimately accomplished by the binding of ammonium (NH_4_ ^+^) to organic acids, biochemical processes to utilize nitrate (NO_3_ ^−^) are of particular importance in aquatic environments because, with the exception of some eutrophic environments, many of the reactive forms of inorganic nitrogen available to autotrophs are restricted to nitrate (Gruber 2008). Therefore, most photoautotrophs possess a two-step nitrate assimilation pathway in which nitrate is reduced to ammonium for utilization. Generally, in plants and algae, nitrate reductase in the cytosol first reduces nitrate to nitrite (NO_2_^−^), and nitrite reductase in the chloroplast stroma reduces nitrite to ammonium (Guerrero et al. 1981, Fernandez and Galvan 2008, Sanz-Luque et al. 2015). In contrast, heterotrophs are able to utilize organic nitrogen and do not require the active utilization of inorganic nitrogen; indeed, they lack the set of genes for the nitrogen assimilation pathway.

The present study investigated the utilization of inorganic nitrogen by *Rapaza viridis* (Yamaguchi et al. 2012, Karnkowska et al. 2022), a phagotrophic euglenid that exhibits kleptoplasty. The kleptoplasty of *R. viridis* confers phototrophic physiology to this flagellate derived from a heterotrophic lineage. The chloroplast donor (the green alga *Tetraselmis* sp.) is first ingested into the phagosome of *R. viridis*, and then the nucleus and cytoplasm of *Tetraselmis* other than the chloroplast are removed, with finally only the chloroplast remains in the cytoplasm of *R. viridis*. The thus-obtained kleptoplast is exploited as if it were its own actual organelle for photosynthesis. Subsequently, ^13^C isotope labeling experiments showed that inorganic carbon was taken up and incorporated into the polysaccharide grains formed in the cytosol of *R. viridis*, which accumulate only in the light phase and disappear in the dark phase under light–dark cycles during culture (Karnkowska et al. 2022). The cells of *R. viridis* multiply by repeated cell division for approximately one week during the period following kleptoplast acquisition. However, this cell division was observed only in medium containing inorganic nitrogen species such as ammonium and/or nitrate ions. This suggests that, in addition to the supply of organic carbon by photosynthesis, the assimilative use of inorganic nitrogen is essential for *R. viridis*. Indeed, transcriptomic and draft genomic data for *R. viridis* have revealed the presence of a nitrate reductase-like gene (Rv*NaRL*) which is not normally present in heterotrophs. To verify the function of *Rv*NaRL, we performed gene knockdown experiments using RNAi and gene knockout experiments using the CRISPR-Cas9 system. In addition, a phylogenetic analysis was conducted to explore the origin of this atypical gene among heterotrophs.

## Results

### Identification and phylogenetic analysis of putative cytosolic nitrate reductase in Rapaza viridis

A local TBLASTN search was performed against our in-house *Rapaza viridis* RNA-seq database using the amino acid sequence of *Chlamydomonas reinhardtii* as a query; only a single isozyme (named Rv*NaRL*), presumed to be a nitrate reductase, was identified. The whole length of Rv*NaRL* cDNA was amplified by 5’ -RACE and 3’-RACE using primers listed in Table 1 and sequenced (accession number: LC738858). The amplification by 5’-RACE with the typical euglenid spliced leader sequence as the forward primer (Euglenid-SL-Fw in Table 1, Nakazawa et al. 2015) confirmed that the mRNA was transcribed from the genome of *R. viridis* and not from the chloroplast donor *Tetraselmis*. In the TargetP analysis, no targeting signal was predicted in the translated peptide sequence at the N-terminus (Almagro Armenteros et al. 2019), suggesting that the expression of the protein product likely occurred in the cytosol. The corresponding sequence was amplified from the genomic DNA of *R. viridis*, revealing that no introns were observed within the transcribed region.

**Table 1.**
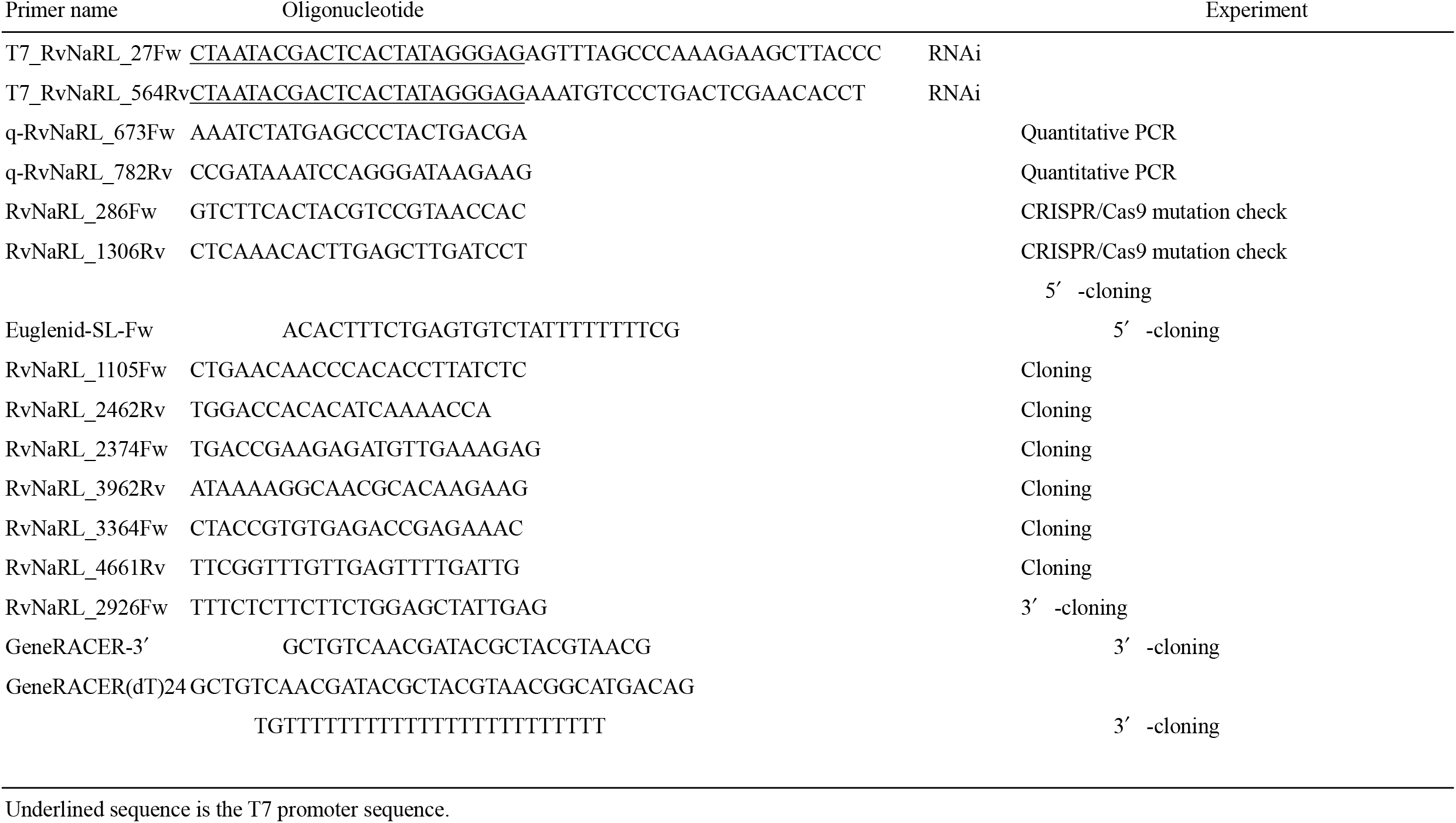
Oligonucleotide primers used in this study.

Phylogenetic analysis, as inferred from Rv*NaRL* and previously reported eukaryotic nitrate reductase homologs, failed to demonstrate the clear relationship of Rv*NaRL* to any homolog of other taxa (Fig. 1). Most notably, a nitrate reductase of Euglenophyceae was shown in the single clade with Rhodophyceae from which Rv*NaRL* was excluded; nonetheless, Euglenophyceae was the closest sister lineage of *R. viridis* in the phylogeny based on the 18S rDNA sequence (Yamaguchi et al. 2012). However, the overall low bootstrap values of the nodes near the base of the phylogenetic tree did not provide a clear indication of the origin of *Rv*NaRL.

**Fig. 1.**
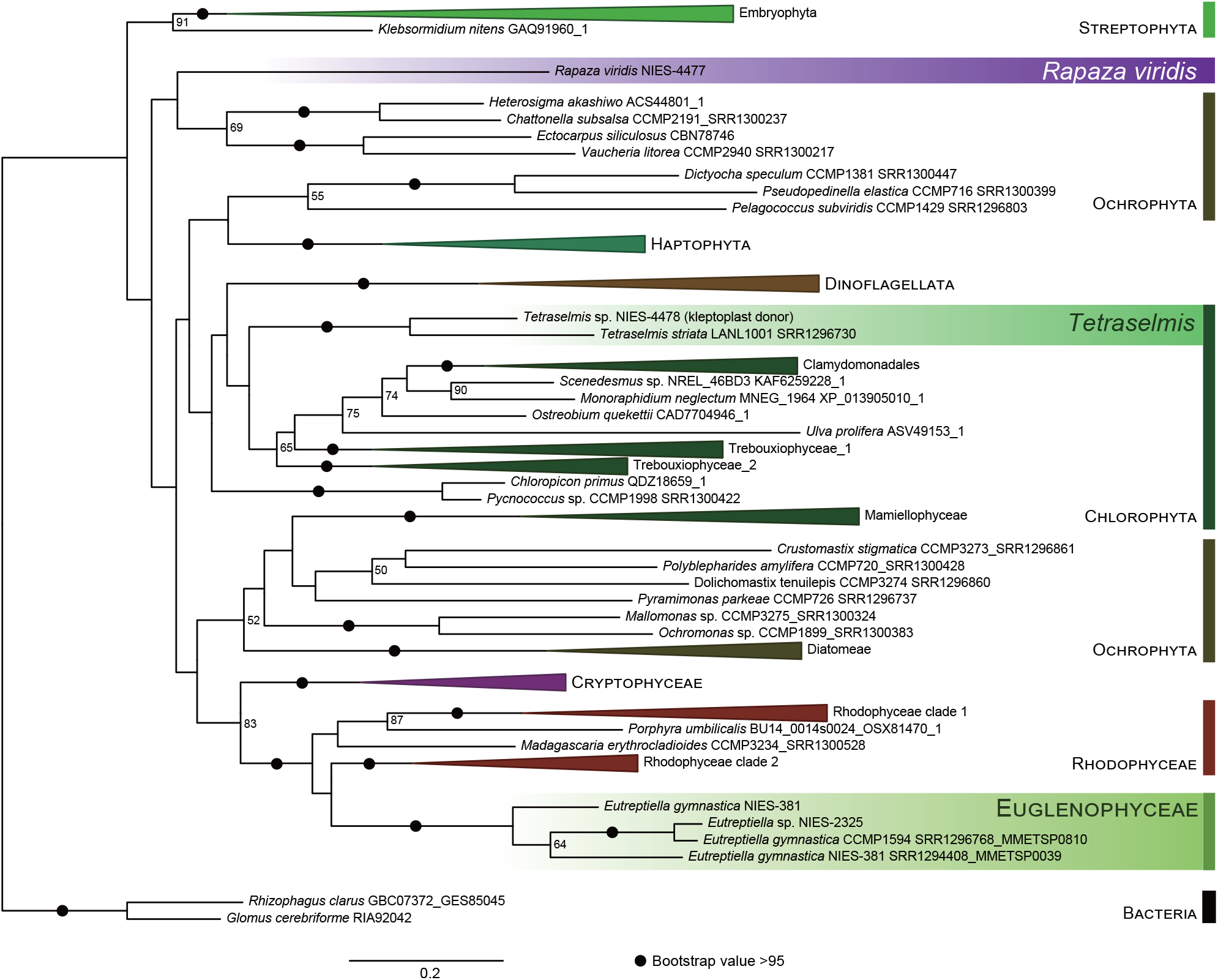
Phylogeny of nitrate reductase suggesting independent evolutionary origins (i.e., independent horizontal gene transfer events) between the *Rapaza viridis* homolog and those of the sister phototrophic clade Euglenophyceae. Maximum likelihood tree of the NaR estimated by 100 rapid bootstrapping replicates in IQ-TREE. Black circles (•) denote bootstrap support >95; support values below 50 are not shown.

### RvNaRL knockdown and knockout experiments in Rapaza viridis

To investigate the function of the gene product *Rv*NaRL *in vivo*, we performed RNAi-mediated gene silencing experiments by introducing double-strand RNA containing a partial Rv*NaRL* sequence. To verify the degree of gene silencing during the experiments, cell aliquots were collected for RNA extraction at each experiment/observation for quantitative RT-PCR, which was performed to confirm that the mRNA expression was maintained at a significantly lower level compared with the wild-type strain during the culture experiment (Supplementary Fig. S1). Hereafter, RNAi-mediated *Rv*NaRL knockdown cells are referred to as Rv*NaRL*-kd cells.

In parallel, Rv*NaRL* knockout lines were prepared by gene editing using CRISPR/Cas9, in which a guide RNA was designed at the position corresponding to the N-terminus region of *Rv*NARL protein to introduce an indel that causes a frameshift owing to a non-homologous end-joining error after Cas9 nuclease activity (Fig. 2A). Whole primary cultures were extracted one week after the introduction of ribonucleotide protein (RNP; the complex of Cas9 and guide RNA), and the PCR-amplified products of the region spanning the edited site were analyzed using SeqScreener Gene Edit Confirmation App (https://apps.thermofisher.com/apps/gea-web/#/setup), resulting in a mutation rate of 68.6% and a frame shift rate of 56.0%. Then, three clones were selected and isolated from the primary culture, and sequencing was performed for the gene editing site of each of three clones exhibiting a single nucleotide deletion (strain *del*1), a single nucleotide insertion (strain *ins*1), and an eleven-nucleotide insertion (strain *ins*11), respectively (Fig. 2B), to identify the downstream frameshifts and stop codons (Fig. 2C). Hereafter, the cells of the three mutant strains are referred to as Rv*NaRL*-KO cells. Cells in the three knockout lines exhibited indistinguishable phenotypes and were each represented by three replicate *RvNaRL*-KO cells in the subsequent culture experiments (n = 3).

**Fig. 2.**
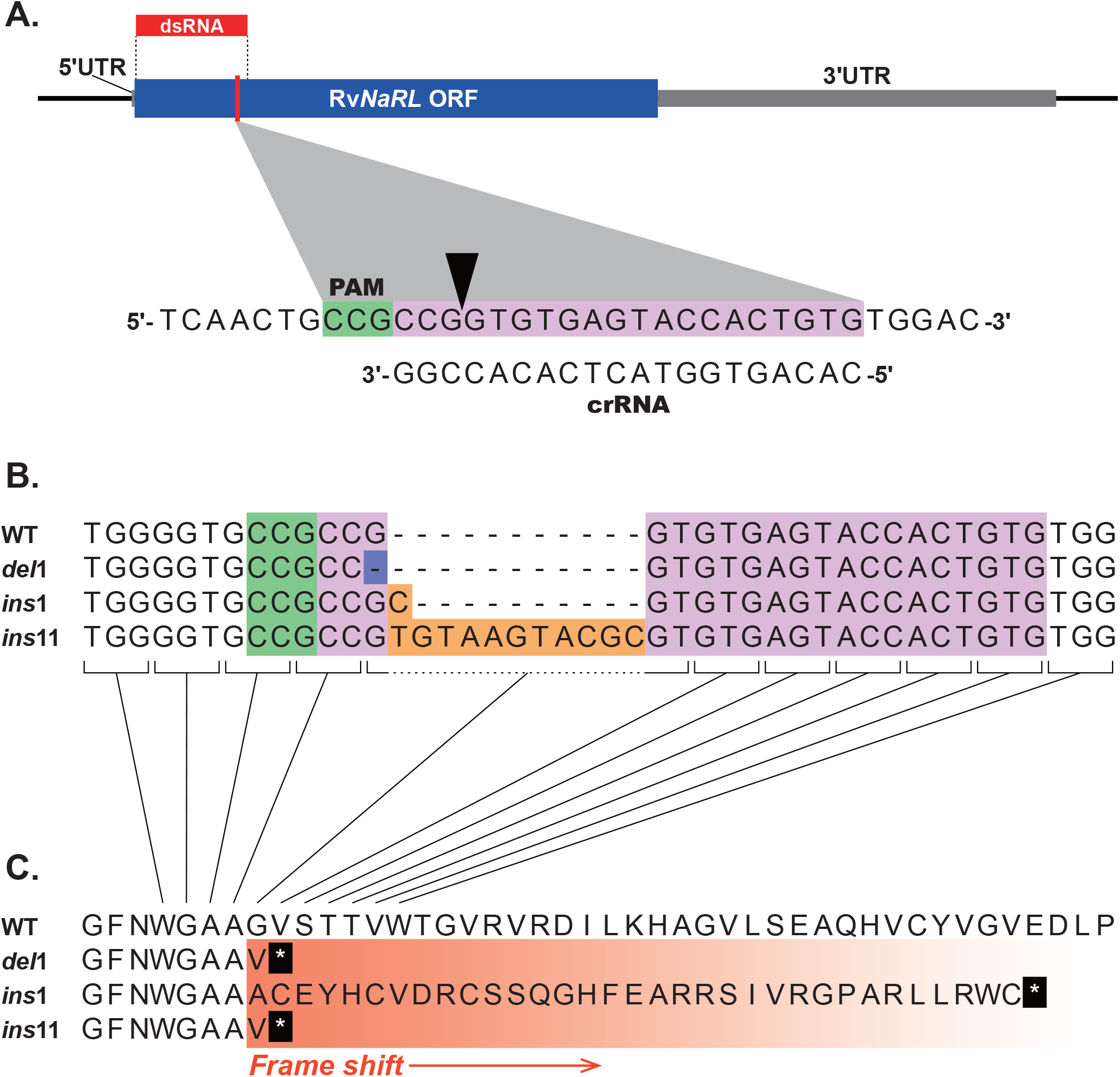
Targeted disruption of the Rv*NaRL* gene. A, Schematic of the Rv*NaRL* gene showing the target (highlighted in pink) and reverse complement PAM (highlighted in green) sequences. The arrowhead indicates the Cas9 cleavage site. B, Partial DNA sequences of the wild-type and three Rv*NaRL* knockout clones. The changes in the target sites compared to the wild-type are shown; the deleted base pair is shown in blue and the inserted base pairs are shown in orange. C, Partial transcribed amino acids sequences of the wild-type and the the Rv*NaRL* knockout clones. Asterisks indicate termination of transcription by a stop codon in the knockout clones.

### Culture experiments with different inorganic nitrogen resources

Rv*NaRL*-kd and Rv*NaRL*-KO cells, as well as cells receiving the blank electroporation treatment, in the RNAi experiments (hereafter referred to as blank cells) and wild-type cells (used as a control for the Rv*NaRL*-KO lines) were cultured in medium containing nitrate as the sole source of nitrogen (nitrate medium) and a medium containing no nitrogen source (N-free medium) as a nitrogen control. The results demonstrated that *RvNaRL*-kd and *RvNaRL*-KO cells did not display active cell growth in the nitrate medium culture and that the cell densities remained at low levels compared with the blank cells and the wild-type cells. Such suppression of growth was comparable to those in the N-free medium for the Rv*NaRL*-kd, Rv*NaRL*-KO, and wild-type cells. Namely, the cell density of the blank cells and the wild-type cells increased 22-fold and 23-fold from the initial state, respectively, suggesting that the cells underwent approximately five cell divisions during this period. In contrast, the cell densities of the *RvNaRL*-kd and *RvNaRL*-KO cells in the nitrate medium, as well as those in the N-free medium, respectively, increased only by approximately 4-fold and 5-fold from the initial state, respectively, suggesting these cells underwent approximately two cell divisions during this period (Fig. 3A).

**Fig. 3.**
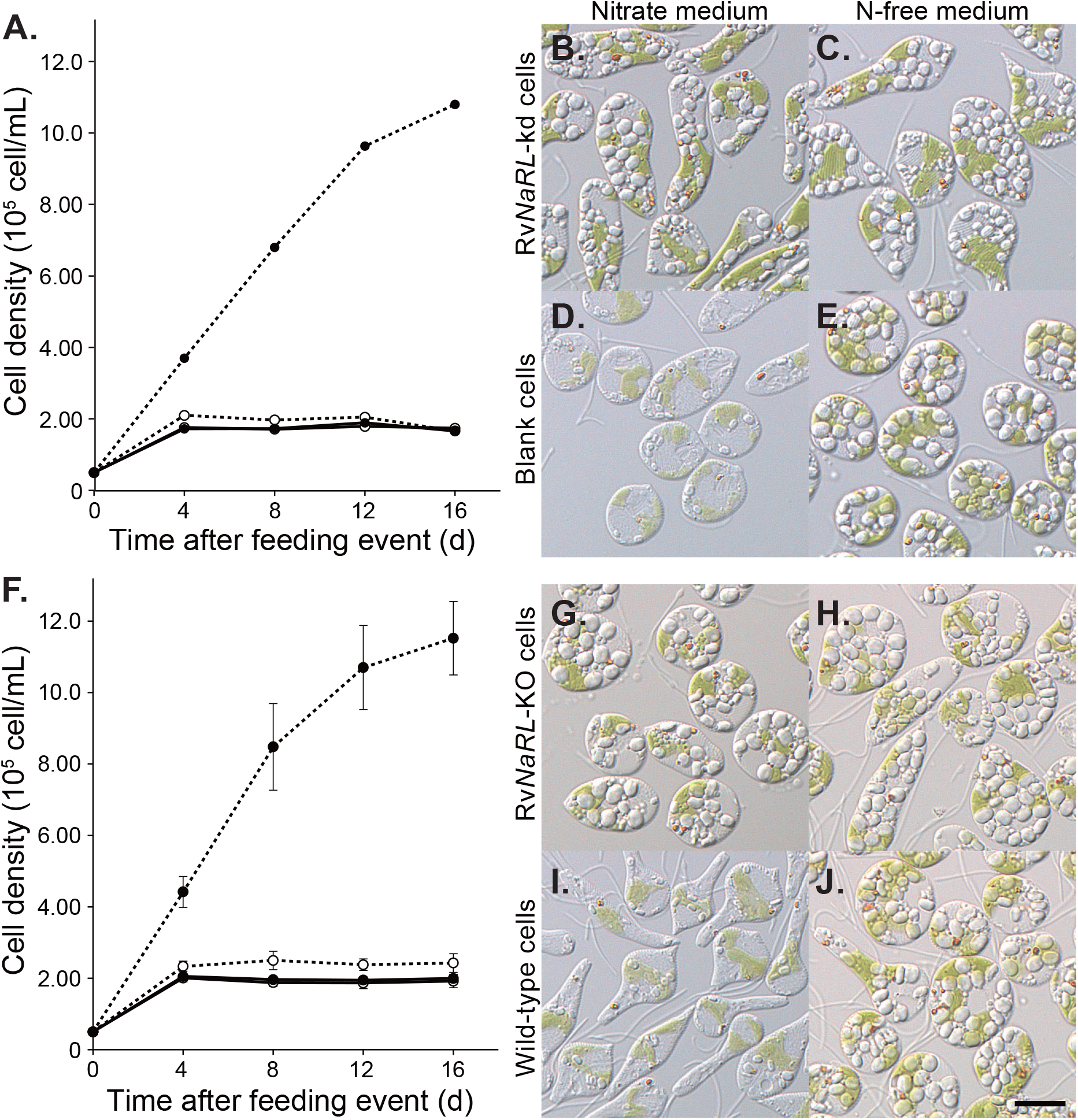
Phenotypes of Rv*NaRL* knockdown and knockout cells grown under ammonium-free culture conditions (nitrogen source: either nitrate or absent). A, growth curves of Rv*NaRL*-kd cells and cells from the blank electroporation experiment (blank cells) in batch culture with each medium. Solid lines represent the growth curves of Rv*NaRL*-kd cells, dashed lines represent those of wild-type cells, • represents nitrate medium, and ○ represents N-free medium. Significant growth was observed only in blank cells cultured in nitrate medium, with almost identical growth inhibition patterns in the other experiments. Day 0 is the day when *Tetraselmis* sp. was fed on the day after the second electroporation experiment. Differential interference contrast (DIC) images on Day 8 of the culture of Rv*NaRL*-kd cells in nitrate medium (B) and N-free medium (C), and of blank cells cultured in nitrate medium (D) and N-free medium (E); the results show similar levels of significant accumulations of polysaccharide grains, except for the blank cells in grown nitrate medium. F, growth curves of the Rv*NaRL*-KO and wild-type cells in batch culture with each medium. Solid lines represent the average growth curves of Rv*NaRL*-KO cells (three independent experiments with different clones: *del*1, *ins*1, and *ins*11 in Fig. 2), dashed lines represent those of wild-type cells (triplicate experiments), • represents Nitrate medium, and ○ represents N-free medium, respectively. As in the knockdown experiments, significant growth was observed only in wild-type cells cultured in nitrate medium, with almost identical growth inhibition patterns in the other experiments. Day 0 is the day when *Tetraselmis* sp. was fed. DIC images on Day 8 of the Rv*NaRL*-kd cells cultured in nitrate medium (G) and N-free medium (H), and of wild-type cells cultured in nitrate medium (I) and N-free medium (J); the results show similar levels of significant accumulations of polysaccharide grains, except for the wild-type cells grown in nitrate medium. All scale bars: 10 μm.

*RvNaRL*-kd, *RvNaRL*-KO, and wild-type cells exhibited significant accumulations of polysaccharide grains in the cytosol when cultured in the N-free medium (Fig. 3B, 3C, 3E, 3G, 3H, and 3J). Both *RvNaRL*-kd and *RvNaRL*-KO cells exhibited comparable accumulations of the polysaccharide grains in the nitrate medium. However, in contrast, only the blank cells and the wild-type cells in the nitrate medium exhibited greatly reduced accumulation of polysaccharides, in which considerably fewer grains with smaller grain size were observed (Fig. 3D and 3I).

Such suppression of growth and the accelerated accumulation of the polysaccharide grains in the knockout cells did not occur when ammonium was present in the medium. Here, to avoid ammonium toxicity to *R. viridis*, only 1/4 molar equivalent of ammonium was added to the nitrate in the nitrate medium as the sole nitrogen source (ammonium medium). Accordingly, medium with a 75% reduction in nitrate concentration was used in the control experiment (1/4 nitrate medium). Almost identical growth patterns and cellular phenotypes were observed for the Rv*NaRL*-KO cells and the wild-type cells in the ammonium medium, suggesting that genome editing had no effect on ammonium utilization in *R. viridis*. These growth rates were somewhat lower than in wild-type cells in the 1/4 nitrate medium (ca. 70%) (Fig. 4A). This could be due to the ammonium toxicity, because the growth was further suppressed when cultured at higher ammonium concentrations (data not shown). Both the Rv*NaRL*-KO cells and the wild-type cells accumulated very few polysaccharide grains in the ammonium medium (Fig. 4B and 4D). In contrast, the RvNaRL-KO cells significantly accumulated polysaccharide grains in 1/4 nitrate medium (Fig. 4C), whereas only a moderate degree of accumulation occurred in the wild-type cells (Fig. 4E).

**Fig. 4.**
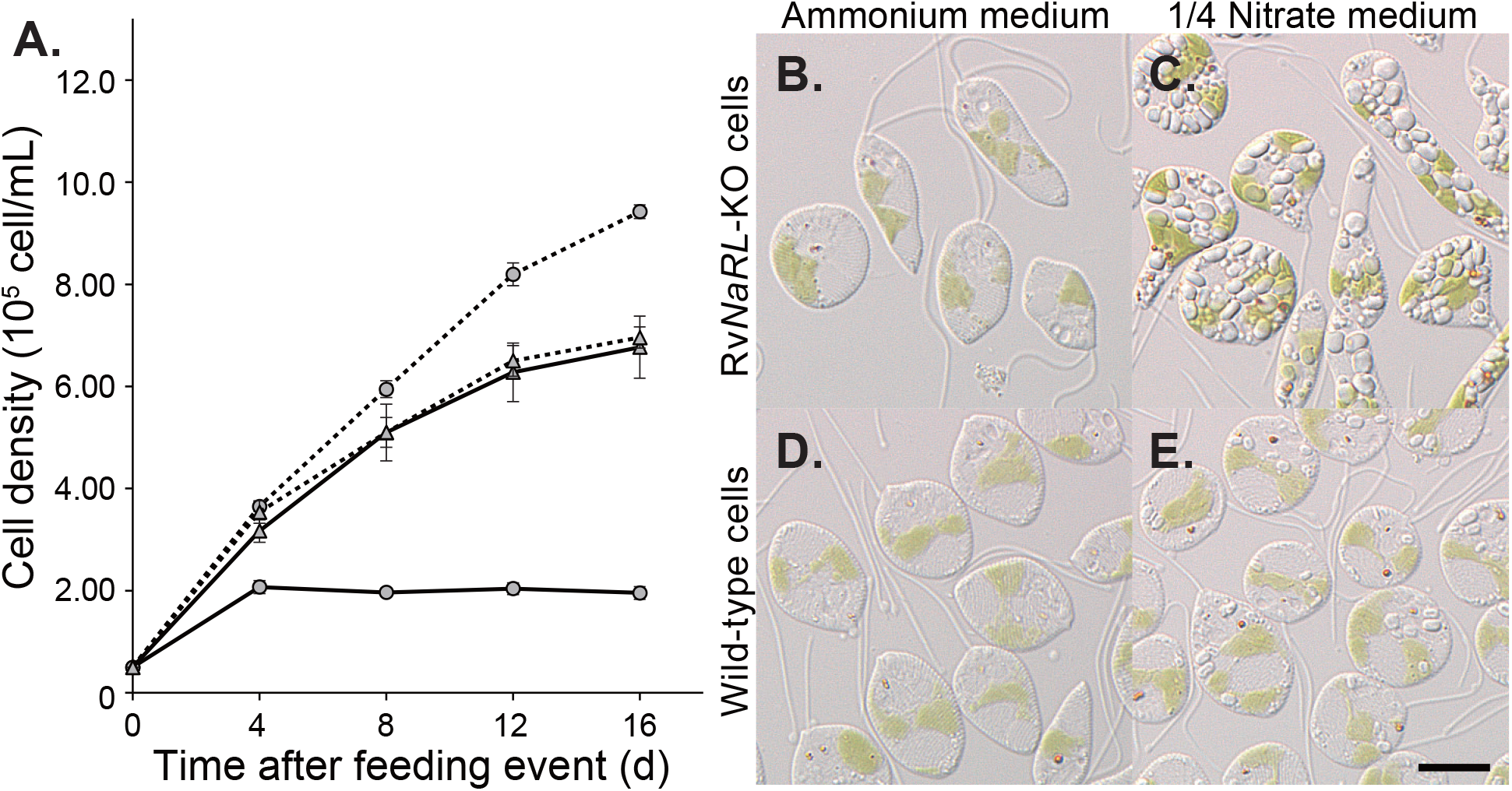
Comparison of phenotypes in cultures with ammonium or nitrate as the sole nitrogen source for the Rv*NaRL*-KO and wild-type cells, respectively. A, growth curves of the Rv*NaRL*-KO cells and wild-type cells in batch culture with each medium. Solid lines represent the average growth curves of the Rv*NaRL*-KO cells (three independent experiments with different clones: *del*1, *ins*1, and *ins*11 in Fig. 2), dashed lines represent those of wild-type cells (triplicate experiments), ▴ represent ammonium medium, and • represents 1/4 nitrate medium. Both the Rv*NaRL*-KO and wild-type cells displayed similarly significant growth curves in ammonium medium, which was a little less than that of the wild-type cells in 1/4 nitrate medium. Day 0 is the day when *Tetraselmis* sp. was fed. DIC images of the Rv*NaRL*-kd cells cultured in ammonium medium (B) and 1/4 nitrate medium (C), and wild-type cells cultured in ammonium medium (D) and 1/4 nitrate medium (E), among which little accumulation of polysaccharide grain was observed in ammonium medium for both the Rv*NaRL*-KO and wild-type cells. All scale bars: 10 μm.

## Discussion

*Rapaza viridis* depends the photosynthesis of the kleptoplast to obtain organic carbon required for its biological activity (Karnkowska et al. 2022). Somewhat unusually, *R. viridis* does not appear to digest the cytoplasm of *Tetraselmis* sp., a chloroplast donor ingested by phagocytosis; moreover, it does not exhibit any other heterotrophic activity (no phagocytosis of other objects or osmotrophic growth). With respect to nitrogen utilization, our present results also strongly suggest that this flagellate uses its own enzymes to assimilate inorganic nitrogen rather than organic nitrogen derived from the chloroplast donor, confirming the above observations. Consequently, the wild-type cells of *R. viridis* did not increase much in number in the absence of inorganic nitrogen in the medium, whereas cell growth was observed only in the medium containing nitrate or ammonium (Fig. 3F and 4A). Under such inorganic nitrogen deprivation, the accumulation of cytosolic polysaccharide grains was accelerated in *R. viridis* (Fig. 3J), a phenomenon similar to that observed in *Euglena gracilis* under nitrogen-deprived conditions as a consequence of the impeded amino acid synthesis, which results in direct transport of the carbohydrate supplied by photosynthesis into the cytosol and its accumulation therein (Coleman et al. 1988).

Regarding nitrate assimilation in particular, our results indicated that *R. viridis* achieved this metabolism, atypical in heterotrophic lineages, by the action of the nitrate reductase encoded in its own nuclear genome. In cells in which Rv*NaRL* was knocked down or knocked out, cell growth was largely suppressed in the presence of nitrate, as observed in wild-type cells in the complete absence of inorganic nitrogen (Figs. 3A and 3F), and the accumulation of polysaccharide grains in the cytosol was significantly enhanced (Figs. 3B and 3G). It can therefore be concluded that the translated product *Rv*NaRL actually functions as a unique enzyme responsible for nitrate reduction in *R. viridis*. The nitrite produced by this enzyme is expected to be further reduced to ammonium by either the residual nitrite reductase of *Tetraselmis* sp. in the kleptoplast or by an enzyme unique to *R. viridis*, although a candidate for this has not yet been identified.

It has been reported that dinoflagellates of the genus *Dinophysis* which, like *R. viridis*, exhibits kleptoplasty, do not assimilate nitrate from the culture medium (García-Portela et al. 2020, Hattenrath-lehmann et al. 2021). In the case of *Dinophysis acuminata*, significant growth was observed under conditions in which photosynthesis by the kleptoplasts provided organic carbon and the assimilation of ammonium in the medium provided organic nitrogen (Hattenrath-lehmann et al. 2021). Here, the assimilation of inorganic nitrogen was assumed to be necessary to balance the profit from the kleptoplast (Hattenrath-lehmann et al. 2021). However, no clear evidence for the nitrate reductase gene has been found among *Dinophysis* in which nitrate assimilation has indeed not been observed (García-Portela et al. 2020, Hattenrath-lehmann et al. 2021).

In contrast, in the case of *R. viridis*, the ability to utilize nitrate as an inorganic nitrogen source must be an ecological advantage. Ammonia exerts a certain degree of toxicity on *R. viridis*, and cell growth is significantly inhibited in medium with equal molar amounts of ammonium to that of nitrate in standard marine algal medium. In the real-world ocean surface environment, the only major source of inorganic nitrogen available in most cases is nitrate; ammonium is rarely present in sufficient concentrations as in laboratory culture conditions (Gruber 2008). The nitrate assimilation pathway is therefore very common among marine photoautotrophs, but this present study of *R. viridis* is the first example of its presence being confirmed through both molecular and biochemical experiments among heterotrophic lineages that include kleptoplastic organisms.

One possible hypothesis for the origin of *Rv*NaRL in *R. viridis*, which exhibits kleptoplasty, is horizontal gene transfer from the chloroplast donor *Tetraselmis* sp. However, because there was a lack of strong support for the phylogenetic relationships of nitrate reductase between most of the higher taxa, we were unable to test the proximity of *Rv*NaRL to the NaR of *Tetraselmis*. In contrast, NaR of all Euglenophyceae, a sister lineage group of *R. viridis*, as suggested by phylogenetic analysis of SSU rRNA genes (Yamaguchi et al. 2012), formed a monophyletic clade with a strong link to all Rhodophyta (Fig. 1). This strongly suggests that the former was acquired by horizontal gene transfer from the latter. Interestingly, however, *Rv*NaRL was not included in this clade, so we could conclude that *R. viridis* acquired this gene by a horizontal transfer independent of Euglenophyceae. Indeed, the acquisition of various exogenous genes by horizontal transfers to *R. viridis* was thought to have facilitated the evolution of kleptoplasty in this organism (Karnkowska et al. 2022). The same is likely true for the acquisition of *Rv*NaRL, which was a key factor in allowing *R. viridis* to evolve into a kleptoplasty-based photoautotroph.

Similarly, photosynthetic cells also possess a plasma membrane nitrate transporter for the uptake of nitrate into the cell (Fernandez and Galvan 2008). Naturally, the presence of a nitrate transporter is also expected in *R. viridis*, as nitrate utilization was observed. Various other mechanisms, such as a control mechanism to prevent the accumulation of toxic nitrite, may also be necessary for the correct functioning of the nitrate assimilation pathway (Stitt and Krapp 1999, Galván and Fernández 2001). It remains to be determined which regulatory mechanisms and sets of protein factors are involved in the association with the exotic kleptoplasts in the nitrate assimilation pathway of *R. viridis*; this information will be essential for understanding the kleptoplasty and, ultimately, the evolution that led to the acquisition of chloroplasts.

## Materials and Methods

### Organisms and cultures

Both strains of *Rapaza viridis* (NIES-4477) and *Tetraselmis* sp. (NIES-4478) were axenically maintained in Daigo IMK medium (Nihon Pharmaceutical, Tokyo, Japan; the maintenance medium), which contained 2.4 mM nitrate and 0.050 mM ammonium, incubated at 20°C under illumination (100–150 μmol-photons m^−2^ s^−1^) by a 14 h light/10 h dark photoperiod (14L:10D). Cultures of *R. viridis* were supplied with three times as many cells of the kleptoplast donor *Tetraselmis* sp. (1:3 ratio) upon transfer to fresh medium every 2 weeks; however, the latter were usually completely consumed within 12 hours, so that the former cultures were essentially maintained in a monospecific state (Karnkowska et al. 2022). The experimental media used in the present study were f/2 medium-based compositions containing only nitrate, only ammonium, or neither. The nitrate medium contained 0.88 mM nitrate, the ammonium medium contained 0.22 mM ammonium, the 1/4 nitrate medium contained 0.22 mM nitrate, and the N-free medium contained neither nitrate nor ammonium. Before culture in the experimental medium, *R. viridis* cells were first fed with three times as many chloroplast donor *Tetraselmis* sp. cells in maintenance medium to allow *R. viridis* to engulf all the donor cells in culture. After complete consumption, all cells were pelleted and transferred to the experimental medium. These operations were intended to eliminate the possibility that differences in the experimental medium would affect the time required for the consumption. The cell numbers of both strains were counted using a particle counter/analyzer (CDA-1000, Sysmex, Kobe, Japan).

### Light microscopy

Differential interference contrast images were obtained from an inverted microscope (IX71, Olympus, Tokyo, Japan) equipped with a color CCD camera (FX630, Olympus) and a Flovel Image Filing System (Flovel, Chofu, Japan). The cells were placed between two glass coverslips (0.13–0.17 mm) and observed.

### Cloning of genomic DNA of the region encoding RvNaRL

The nucleotide sequences of genomic DNA of the region encoding Rv*NaRL* was subjected to BLASTN searching in our in-house DNA-seq database, and a single contig therein was contained the entire sequence of the mRNA, except for the 5’ spliced leader and the 3’ poly(A) sequences. The contig sequences were confirmed by PCR amplification and sequencing of genomic DNA using primers designed based on the cDNA sequences (primers not shown).

### Phylogenetic analysis

BLASTP searches were performed against the NCBI nr database and The Marine Microbial Eukaryote Transcriptome Sequencing Project (Keeling et al. 2014) using the translated peptide sequence of *Rv*NaRL as a query. The homologs of the chloroplast donor *Tetraselmis* sp. and the relatives of *R. viridis* host lineage, *Eutreptiella*, were retrieved from in-house RNA-seq data and manually added to the dataset. The sequences were aligned using the MAFFT algorithm with the default parameters by the MAFFT package v7.490 (Katoh et al. 2019) and then trimmed by trimAl 1.2rev57 (Capella-Gutierrez et al. 2009). After the manual exclusion of short and ambiguous aligned positions, 611 amino acid sites and 250 OTUs remained. ML trees were inferred by IQ-TREE software v2.0.3 (Trifinopoulos et al. 2016) with nonparametric bootstrapping (100 replicates) under the LG+R8 model. The tree was inspected manually to examine the relationship between the *R. viridis* sequence and the sequences of other organisms in the database.

### RNAi Experiments

Silencing experiments using RNAi targeting Rv*NaRL* were performed employing the method designed for *E. gracilis* (Iseki et al. 2002, Nakazawa et al. 2015) with modifications. Partial cDNA of the Rv*NaRL* sequence was PCR-amplified with the addition of the T7 RNA polymerase promoter sequence at the 5’-ends of both primers (T7_*Rv*NaRL_27Fw and T7_*Rv*NaRL_564Rv; listed in Table 1) using PrimeSTAR GXL DNA Polymerase, and the subsequent PCR product was purified using Wizard SV Minicolumns (Promega). Double-strand RNAs (dsRNAs) containing a partial Rv*NaRL* sequence (590 bp) were synthesized and purified using the MEGAscript RNAi kit (Thermo Fisher Scientific, Massachusetts, USA) in accordance with the manufacturer’s instructions. At 1 week after *R. viridis* cells were fed *Tetraselmis* sp. In Daigo IMK medium, the cells were collected (4 × 10^6^ cells), washed in electroporation (EP) buffer (500 mM solution of trehalose in 5% seawater) twice, resuspended in 200 μL of the EP buffer containing 15 μg of the dsRNA, and incubated on ice for 90 seconds. The cell suspension was transferred to a 0.2 cm gap cuvette and immediately electroporated by GENE PULSER II (Bio-Rad, Hercules, CA, USA) at 0.45 kV and 50 μF. The cell suspension was then immediately inoculated into fresh medium and incubated at 20°C in the dark for 10 hours before incubation in the 14L:10D environment at 20°C for an additional 36 hours. All cells incubated under these conditions were collected again and treated once more with the electroporation procedure described above to obtain a culture composed predominantly of knockdown cells. In parallel, as a blank test for the dsRNA introduction, a control electroporation experiment, using the same procedure, was performed on cells suspended only in EP Buffer without dsRNA.

### Quantitative PCR

Total RNA was extracted from pelleted Rv*NaRL* knockdown and blank experimental cells using ISOGEN II (NIPPON GENE, Tokyo, Japan), from which cDNA was obtained by reverse transcription using the PrimeScript RT Reagent Kit (TaKaRa, Kusatsu, Japan). Transcript levels of Rv*NaRL* were examined using a real-time PCR system (LightCycler 96 system, NIPPON Genetics, Tokyo, Japan) with KAPA SYBR Fast Universal (NIPPON Genetics) and the primers listed in Table 1.

### CRISPR/Cas9 genome editing and preparation of the knockout clones

A 20-nucleotide CRISPR RNA (crRNA) guide sequence (CACAGUGGUACUCACACCGGCGG) was designed using the CRISPRdirect website (Naito et al. 2015). The crRNA was custom synthesized by Integrated DNA Technologies (IDT) as a chemically modified 36 nt oligo with 16 additional nucleotides annealing to the tracrRNA. Single-guide RNA (sgRNA) was then prepared by mixing 0.6 μL each of the 100 μM crRNA and 100 μM tracrRNA (Alt-R CRISPR-Cas9 tracrRNA, IDT) in a 200 μL PCR tube and incubated at 95°C for 5 minutes before cooling slowly to room temperature for hybridization. Then, 0.8 μL of Guide-it Recombinant Cas9 (10 μg/μL, Takara) was added to the above PCR tube with the prepared sgRNA and incubated at 37°C for 5 min to assemble the ribonucleotide protein (RNP) complex. RNPs designed to cleave approximately 20% of the N-terminal side of the *Rv*NaRL peptide sequence when guided by the above sequence were introduced into *R. viridis* cells by the above electroporation method with the following modifications. *R. viridis* cells (2 × 10^5^ cells) were collected and finally resuspended in the PCR tube, in which 50 μL of the EP buffer was pre-mixed with the RNP prepared above (in total, 70 μL) and cooled. The cell suspension after electroporation was then immediately inoculated to fresh medium and incubated at 26°C in the dark for 24 hours. The RNP-transfected *R. viridis* cells were then fed with an equal number of *Tetraselmis* sp. cells and further cultured in a 14L:10D environment at 26°C for 72 hours before being transferred to a normal culture environment at 20°C.

RNP-transfected *R. viridis* cells were collected 1 week after the transfection and processed for genomic DNA (gDNA) extraction using phenol/chloroform/isoamyl alcohol (25:24:1) (NIPPON GENE). The region containing the edited site was amplified from the extracted gDNA by PCR using KOD -Multi & Epi- (Toyobo, Osaka, Japan) with forward and reverse primers (*Rv*Contig439_2147Fw and *Rv*NaRL_1306Rv, respectively; Table 1) followed by column purification using NucleoSpin^®^ Gel and PCR Clean-up (Macherey-Nagel, Düren, Germany). After sequencing the purified PCR product, the DNA electropherogram file (AB1 file) was uploaded to and analyzed using SeqScreener Gene Edit Confirmation App (https://apps.thermofisher.com/apps/gea-web/#/setup (accessed on November 06, 2021) to assess the frequency of the targeted mutation in the primary culture after transfection. Clonal strains were generated from the primary culture by aseptically separating individual *R. viridis* cells into cultural plates by the microcapillary method using glass tubes under an inverted microscope (CKX-41, Olympus) housed in a laminar flow cabinet. The editing sites were analyzed in the same manner as described above, and the mutated sequences of each clone were determined.

## Acknowledgements

We thank Tomomi Munekyo and Toru Shirasaki for their technical assistance with the molecular biological experimental operations.

## Legends to Figures

**Fig. S1.**
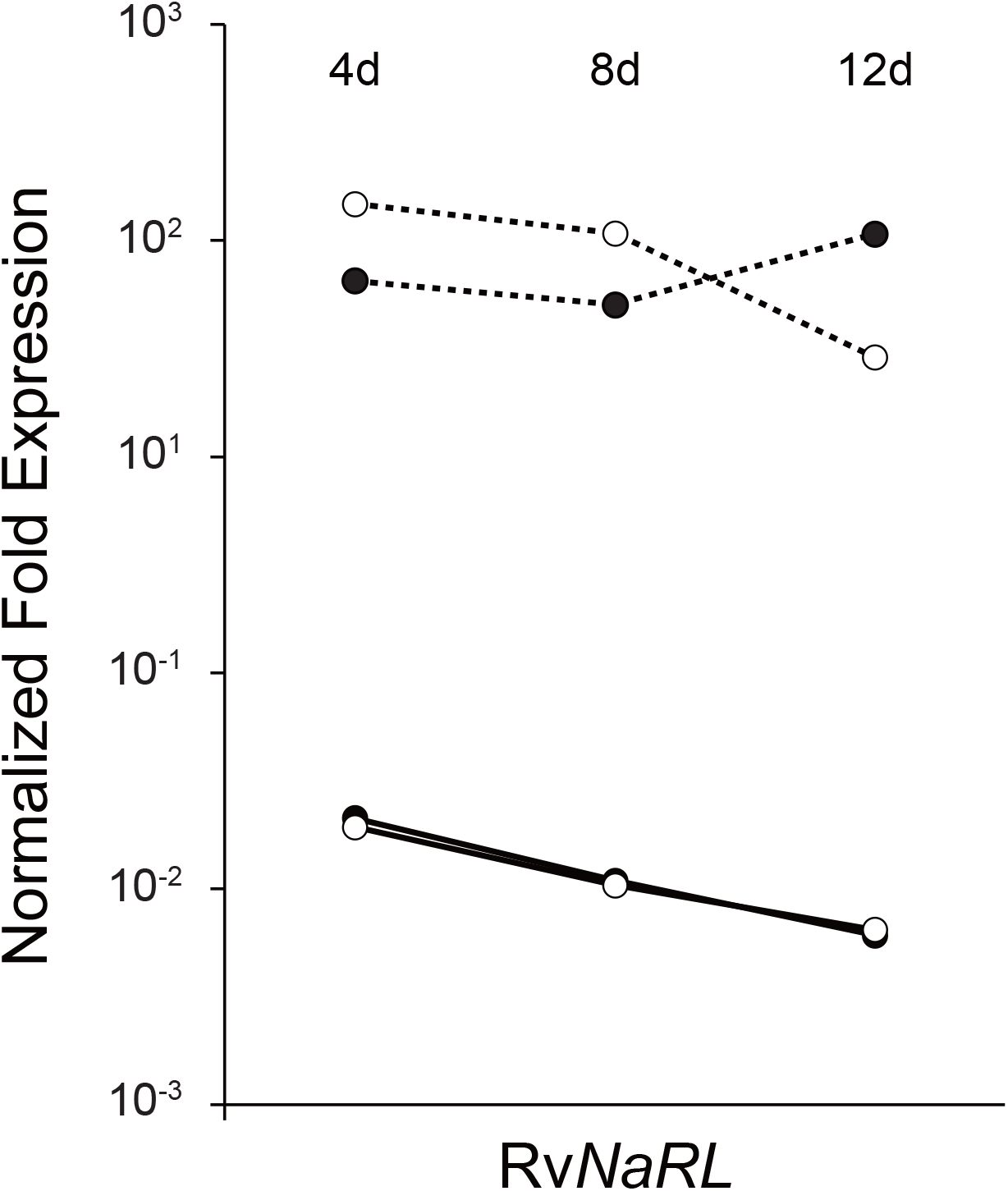
Quantitative evaluation of transcript levels of Rv*NaRL* using qPCR. Values shown are normalized to α-tubulin transcript levels. The solid line represents the Rv*NaRL*-kd cells, the dashed line represents the blank cells (electroporation experimental control cells), • represents nitrate medium, and ○ represents N-free medium. Rv*NaRL* expression in the whole culture of Rv*NaRL*-kd cells is approximately 1/1000 lower than that in the blank cells.

